# Loss of Vitellogenin Receptor Function Results in Yolk Depletion, Virome Expansion and Reduced Bacterial Load Within the Oocytes of *Rhodnius prolixus*

**DOI:** 10.64898/2025.12.05.692514

**Authors:** Juliana Amorim, Jéssica Pereira, Matheus das Neves, Thamara Rios, Cintia Lopes Nogueira, Valdir Braz, Leonan Reis, Allana Faria-Reis, Ana Beatriz Walter-Nuno, Lukas Selim, Daiene Nunes, Thalysson Vinícius de J. C. Baptista, Georgia Atella, Marinella Silva Laport, Ana C. Bahia, Carlos Logullo, Pedro L. Oliveira, Gabriela O. Paiva-Silva, Katia C. Gondim, Isabela Ramos

## Abstract

The vitellogenin receptor (VgR) mediates yolk protein uptake during oogenesis and is essential for embryogenesis in oviparous species. Here we characterize the single *Rhodnius prolixus* VgR isoform and uncover an unexpected role in microbial regulation within the reproductive system. The receptor displays a conserved LDLR-like structure and is highly expressed in early oocytes. RNAi-mediated VgR silencing caused defective yolk granule biogenesis, leading to the accumulation of the main yolk protein precursors, Vg and RHBP, in the hemolymph, yet oviposition and fertilization proceeded normally. The resulting eggs were yolk-depleted and non-viable. Remarkably, VgR knockdown reduced bacterial 16S rRNA levels in oocytes while promoting the expansion of several members of the core virome, a phenotype not reproduced by Vg silencing. Neither purified Vg nor changes in immune (defensin) or RNA interference pathways explained the microbial shifts. These findings indicate that VgR governs not only yolk endocytosis but also the trafficking of microbial components into developing oocytes. We propose that VgR contributes to the linking of yolk endocytic dynamics and microbial homeostasis, influencing the balance of microbial components within developing oocytes. This connection broadens the functional scope of the VgR and provides new insight into how vertical transmission processes are shaped in this major Chagas disease vector.

**Author summary:** Egg-laying animals must load their eggs with enough nutrients to support early development. In insects, this process depends on a receptor that brings yolk proteins into the growing egg. Here, we studied this receptor in *Rhodnius prolixus*, a major vector of Chagas disease, and uncovered an unexpected link between yolk uptake and the microorganisms that enter the egg.

When we blocked the receptor, females continued to produce and lay eggs, but these eggs failed to accumulate yolk and could not support embryonic development. Strikingly, the absence of the receptor also shifted the microbial community inside the oocyte: bacterial levels dropped, while several viruses expanded. These changes did not result from differences in yolk proteins, immune activation, or direct antimicrobial effects, indicating that the receptor itself influences microbial entry or persistence in the egg.

Our findings reveal that this yolk receptor plays a dual role, providing nutrients and shaping the microbial community that is passed from mother to offspring. This work highlights an unrecognized layer of interaction between reproduction and microbial transmission in an important disease vector, offering new perspectives for understanding and potentially disrupting vertical transmission pathways.

## Introduction

*Rhodnius prolixus* is a hematophagous hemipteran that serves as one of the principal vectors of *Trypanosoma cruzi*, the causative agent of Chagas disease. This neglected tropical disease (NTD) is endemic to Central and South America, affecting approximately 8 million individuals, with significant morbidity [1]. Additionally, *R. prolixus* holds historical importance as a foundational model organism in studies of insect physiology, endocrinology, and development, with its utility enhanced by the availability of its genome [2]. Consequently, *R. prolixus* remains a dual-focus species of both medical significance and scientific inquiry, enabling advances in molecular and physiological research with implications for vector management [3].

To assure a sufficient supply of nucleic acids, proteins, lipids, phosphate, carbohydrates, ions, and vitamins required for the future embryo’s autonomous growth, developing oocytes in oviparous species, including insects, rely on the accumulation of large volumes of yolk. In most insects, the majority of the components of the yolk are produced extraovarially in the fat body, secreted into the hemolymph and carried into the developing oocytes by membrane-bound receptors via receptor-mediated endocytosis [4,5]. Vitellogenin (Vg) is by far the most prevalent yolk protein in insect species [6]. Vg Receptors (VgRs) are members of the low-density lipoprotein receptor (LDLR) family and mediate the accumulation of Vgs into the oocytes of many insect and vertebrate species [7]. Following binding, Vg/VgR complexes follow a canonical endocytic route leading to the formation of early and late endosomes, with recycling of VgR to the oocytés membrane. The endocytic route ends in the formation of mature membrane-bound organelles, named yolk granules (YG), where Vg is further processed and stored as Vitellin (Vt) for later use [4,8]. Typically, Vt and its products of degradation are the main source of nutrients that supply the embryo with fundamental molecules for *de novo* synthesis until the newly hatched first instar nymphs become capable of feeding [9].

Insect VgRs are distinct due to possessing two clusters of LBD (Ligand-Binding Domain) and EGFD (Epidermal Growth Factor-like Domain), hence forming a different subclass within the LDLR family of receptors. The variations in the quantity and configuration of these structural units are believed to account for the recognition of the many ligands that are associated with this receptor family [7].

Across insects, VgR mRNA is generally expressed throughout oogenesis often peaking in previtellogenic or vitellogenic stages. *In situ* and immunolocalization studies show that VgR protein is evenly distributed in the oocyte during previtellogenesis and later accumulates at the oocyte cortex during vitellogenesis, consistent with its role in mediating yolk uptake [10–13]. Functional studies across diverse arthropod species have consistently demonstrated that RNAi-mediated silencing of the *VgR* invariably disrupts oogenesis and female fertility. Collectively, these findings underscore the conserved and essential role of VgR in insect reproduction [14–22].

Insects host a wide variety of microbial partners, ranging from mutualistic bacteria to pathogenic and symbiotic viruses, many of which have evolved mechanisms to persist and be transmitted across generations. Bacterial endosymbionts such as *Wolbachia*, *Rickettsia*, and *Spiroplasma* are ubiquitous among arthropod taxa and can profoundly influence host reproduction, immunity, and vector competence through vertical transmission via the germ line [23]. The triatomine *R. prolixus*, for instance, harbor the actinobacterium *Rhodococcus rhodnii* in its intestinal tract, a symbiont that provides essential nutrients for the insect’s development. This association is highly species-specific and critical for normal growth and fecundity of the host [24]. Similarly, vertically transmitted viruses are increasingly recognized as widespread and ecologically relevant components of insect microbiomes. Sigma viruses in *Drosophila* species, for instance, are inherited through both eggs and sperm and exemplify long-term host–virus coevolution [25]. Several plant and animal viruses exploit reproductive tissues for transmission. In the brown planthopper *Laodelphax striatellus*, the Rice stripe virus hijacks the VgR pathway to enter ovaries and achieve transovarial transmission [26,27]. These intimate associations illustrate how reproductive molecular machinery can serve as a gateway for microbial persistence in insect populations. Consistent with this, Brito et al. (2021) [28] recently described a complex core virome in *Rhodnius prolixus* oocytes, composed of seven RNA viruses (RpV1–7) that are vertically transmitted to the progeny, unveiling a stable and heritable viral community integrated into the reproductive biology of this major Chagas disease vector.

In this study, we characterized the single *VgR* isoform of *Rhodnius prolixus* and investigated its roles in reproduction, yolk accumulation, and microbial transfer to the oocytes. We show that the *R. prolixus VgR* is a structurally conserved member of the LDL receptor superfamily and is predominantly expressed in the ovary. Functional silencing of *VgR* by RNAi did not affect blood digestion, reproductive endocrine signaling, or oviposition but resulted in defective yolk uptake, leading to the production of yolk-depleted eggs unable to sustain embryogenesis. These phenotypes were accompanied by the accumulation of the main yolk protein precursors (Vg and RHBP) in the hemolymph and by altered bacterial and viral loads in the oocytes, indicating that VgR-mediated endocytosis influences not only yolk deposition but also the transfer of microbial components to the developing oocytes. Finally, *VgR* expression was also detected in male tissues, particularly in the testis, although its silencing did not affect male physiology, fertility, or progeny viability. Together, our findings identify *VgR* as a key regulator of oocyte maturation in *R. prolixus* and reveal its potential involvement in the transovarial passage of microorganisms, highlighting a connection between yolk accumulation and vertical transmission of microbiota in triatomine insects.

## Methods

### Insects and eggs

Adult insects and eggs were kept in a controlled insectarium, with a 12-hour light/dark photoperiod, temperature of approximately 28°C, and relative humidity around 70-80%. As adults, the insects were fed every 21 days with rabbit blood. All procedures follow a protocol approved by the Ethics Committee on the Use of Animals (CEUA-UFRJ), registered under CEUA #123/22.

### Bioinformatics

The VgR sequence of *R. prolixus* was obtained through the transcriptome of several tissues involved in egg production [29] and validated using the genome and transcriptome deposited in the VectorBase database (Rpro C3.2) (www.vectorbase.org). Conserved domains were predicted using the online software SMART (https://smart.embl-heidelberg.de/).

### Total RNA extraction and cDNA synthesis

For total RNA extraction, the midgut (MG), fat body (FB), and reproductive organs (ovary or testis - OV or TEST) were dissected 7 days after blood feeding from adult females and males. All samples were homogenized with the assistance of a plastic potter in Trizol reagent, and the RNA was extracted, measured by spectrophotometry in NanoDrop (Thermo Scientific) at 260 nm and treated with DNase I (Invitrogen). Following, the treated RNAs were used in reverse transcription reactions using the High-Capacity cDNA Reverse Transcription Kit (Applied Biosystems), according to the manufacturer’s protocol.

### PCR/qPCR

PCR reactions used *Rhodnius prolixus* VgR-specific primers previously described (Faria-Reis et al., 2023) to amplify 216 base pair fragments using the following cycling: 10 min at 95°C, followed by 35 cycles of 30 s at 95°C, 30 s at 52°C and 1 min at 72°C and a final extension of 15 min at 72°C. qPCR reactions were carried out in a StepOne RealTime PCR System thermocycler (Applied Biosystems), using SYBR Green PCR Master Kit (Applied Biosystems) and the following parameters: 95°C for 10 min, 40 cycles at 95°C for 15 s and 60°C for 1 min. The relative expressions were calculated using the delta Ct (cycle threshold) obtained using the endogenous genes 18s (RPRC017412) or EF1 (RPRC015041) and expressed as 2^-dCt^ or 2^-ddCt^, depending on the experiment [30]. Under our experimental conditions, both endogenous genes showed stable expression, which supported their use as reference genes in accordance with the MIQE guidelines [31]. The sequences of all primers are described in Table S1.

### Gene silencing via RNAi

Double-stranded RNAs (dsRNAs) were synthesized by the MEGAscript RNAi Kit (Ambion Inc.), using primers designed for specific sequence amplification with the T7 promoter. The dsRNA was designed to generate amplicons of 707 base pairs. The *Escherichia coli* MalE gene encodes a maltose-binding protein and was used as a control dsRNA. Using a Hamilton syringe, each insect was injected with 1 µg of control or experimental dsRNA, according to the following protocols: 1) for mating experiments between VgR-silenced males and wild-type virgin females, dsRNA was injected in the males 13 days before blood feeding. Eight days later, silenced males and virgin females are allowed to mate for 5 days before being blood fed; 2) for all other experiments, adult insects were injected 2 days before blood feeding. Silencing efficiency was subsequently confirmed by qPCR 7 days after dsRNA injection for protocol 1 and 7 days after feeding for protocol 2. The sequences of the primers are described in Table S1.

### Gene silencing phenotype evaluation

After dsRNA injection according to protocol 2, fed males and females were transferred to individual vials. For the digestion experiments, the insects were weighed before and after the blood meal and every two days afterward. For oviposition analysis, eggs laid by individual insects were collected on the same days that the animals were weighed, and approximately two weeks later, the nymphs that hatched were counted. Mortality rates were recorded daily.

### Fertilization status

To determine if fertilization was occurring in control and silenced eggs, a PCR was performed targeting a male-specific DNA sequence (chromosome Y) (GenBank: JX559072.1), as previously reported [32] Briefly, freshly laid eggs from control and silenced individuals were collected, and genomic DNA was phenol extracted and precipitated with 100% ethanol and 2M ammonium acetate. Purified genomic DNA samples were used as templates for amplification using specific primers. The fat body of adult male insects was used as a positive control, and eggs laid by non-mated females were used as a negative control. The PCR product was subsequently visualized in a 2% agarose gel.

### Hemolymph extraction and SDS-PAGE

Following dsRNA injections, hemolymphs from females were collected on day 7 after blood feeding. Once extracted, the hemolymph was diluted 2x in 50 mM HEPES buffer, pH 7.4 containing a cocktail of protease inhibitors (aprotinin 0.3 µM, leupeptin 1 µg/µl, pepstatin 1 µg/µl, PMSF 100 µM and EDTA 1 mM) and approximately 8 mg of phenylthiourea. 0.5 µl of each hemolymph sample was loaded in a 10% and 13,5% SDS-PAGE and stained with Coomassie Blue or silver nitrate [33]. Each biological replicate corresponds to the hemolymph of one insect. injected with dsMal and dsVgR and homogenized in 50 μl of 50 mM pH 7.4 HEPES buffer containing protease inhibitors (aprotinin 0.3 μM, leupeptin 1 μg/μl, pepstatin 1 μg/μl, PMSF 100 μM, and EDTA 1 mm). The equivalent of one-tenth of the egg was loaded in a 10% and 15% SDS-PAGE and stained with silver nitrate. For each biological replicate, samples were prepared using a pool of 3 eggs, each laid by a different female.

### Determination of protein content

Hemolymph and egg samples were collected as described above and the total amount of protein levels was measured by the Lowry method (Folin), using 1–7 µg of BSA as a standard [34] in an E-MAX Plus microplate reader (Molecular devices) using SoftMax Pro 7.0.

### Hemolymph RHBP Titration

RHBP quantification in the hemolymph was performed as described by Walter-Nuno et al. [32]. Briefly, hemolymph samples (5 µL) were collected from females and immediately diluted in 500 µL PBS containing 3–13 µg/mL phenylthiourea to prevent melanization. The absorbance of the Soret band was monitored between 350–450 nm, and the value at hemin saturation, determined from a break in the titration curve, was used to calculate the total RHBP concentration based on the molar extinction coefficient of 0.0645 µM.

### Vitellogenin (Vg) purification

Vg was purified from *Rhodnius prolixus* oocytes following the ammonium sulfate fractionation and gel-filtration procedure described by [35], with minor modifications. Briefly, oocytes collected four to six days after a blood meal were homogenized in 20 mM Tris-HCl (pH 7.0) containing 0.15 M NaCl and a cocktail of protease inhibitors. The homogenate was centrifuged at 11,000 × g for 5 min at 4 °C, and the lipid layer and pellet were discarded. Solid ammonium sulfate was added to the supernatant to reach 45% saturation, and the suspension was gently stirred for 20 min at 4 °C. After centrifugation (11,000 × g, 10 min), the pellet was discarded, and the supernatant was brought to 60% saturation. The resulting precipitate was washed twice with 60% ammonium sulfate and re-extracted by resuspending in a 45% saturated solution followed by centrifugation. The supernatant was dialyzed against 0.15 M NaCl, 10 mM Tris-HCl (pH 7.0), and applied to a Sephadex G-200 column (2.5 × 55 cm) equilibrated with the same buffer. Protein content of each fraction was monitored at 280 nm, and the characteristic Vg fractions were pooled and dialyzed against deionized water.

### Bacterial growth inhibition assay

Vg antibacterial activity was tested following the protocol of [36] with minor modifications. The antibacterial activity of purified *Rhodnius prolixus* Vg was evaluated against *Escherichia coli* ATCC 25922 and *Staphylococcus aureus* ATCC 29213. Bacteria were cultured overnight in Luria–Bertani (LB) medium at 37 °C with shaking, then suspended in sterile saline (0,85%) to reach 0,5 MacFarland, diluted at a 1:200 rate in cation-adjusted Mueller-Hinton (CAMH) broth, and distributed into sterile 96-well plates. Purified Vg was serially diluted in CAMH and added to the bacterial suspensions to a final concentration of 12.5–200 µg/ml. Bacterial growth was quantified after 24h at 37 °C with shaking by measuring OD₆₀₀ using a microplate reader.

### Trypanosoma rangeli Infection

*Trypanosoma rangeli* infection assays were conducted following the procedures previously described [37]. Briefly, adult females of *Rhodnius prolixus* were infected by injection into the hemocoel of 2µl of epimastigotes of *T. rangeli* (1 × 10^4^ parasites/mL) in sterile saline, 3 days after the blood feeding. On the third day after infection, the insects were dissected, and the fat body and ovary were used for the RT-qPCRs.

### Vg antiviral activity – Virus propagation and Mayaro plaque assay

The arbovirus MAYV (MAYV BR/SJRP/LPV01/2015 strain) was propagated in *Aedes albopictus* C6/36 cells. The culture supernatant was collected, centrifuged, aliquoted, and stored at -80°C for subsequent use in cell infection experiments. Viral titers were determined by plaque assay using Vero cells (African green monkey kidney). Cells were maintained in DMEM (ThermoFisher) supplemented with 10% FBS, 7.5% sodium bicarbonate, and 1% L-glutamine. They were seeded as monolayers (∼70% confluence) in 24-well plates and incubated at 37°C with 5% CO₂ for approximately 24 hours before the assay. Aliquots of MAYV (100 µL, 5 x 10⁸ PFU/ml) containing either Tris/HCl 10mM pH 7.4 (control) or Vg (400 µg/ml) were added to the respective cell monolayers. Subsequently, 500 µL of DMEM containing 2% FBS and 1% methylcellulose (Sigma-Aldrich) were added to each well. Plates were incubated at 37°C with 5% CO₂ for 44 hours. Cells were then fixed with a 10% formaldehyde solution (Sigma-Aldrich) for 1 hour. Then, it was washed and stained with crystal violet (0.5%) in a 20% methanol solution for 20 min at room temperature. Excess stain was removed by washing with water. used for lipid extraction. The lipid composition was analyzed by thin-layer chromatography (TLC) on silica gel plates (Merck) using two consecutive solvent systems [38]. Plates were stained with copper reagent, and relative lipid composition was determined by densitometry using TotalLab Quant v11 (TotalLab) with background corrections after comparison with commercial lipid standards (Sigma) [39]. A total of 5 eggs were used for each biological replicate.

### Determination of glycogen content

Egg homogenates were prepared with 5 eggs in 100 μl of lysis buffer containing 200 mM sodium acetate buffer, pH 4.8, and 0.001% Triton X-100 supplied with a protease inhibitors cocktail (Sigma #P8340) and were used to determine glycogen content. For 4 h, the experimental samples were incubated at 40 C° in the presence of 20 μl (1U) of amyloglucosidase, except for the controls, which were prepared under the same conditions but without enzyme. The quantification was made using the Glucox 500 kit (Doles reagents), according to the manufacturer’s instructions.

### Light microscopy

Eggs collected 0-24 h after oviposition were gently disrupted using fine tweezers in 2 μl of PBS on a slide to observe the yolk granules (Ygs) suspension as previously reported [40,41]. The samples were observed using a Zeiss Axio Imager D2 microscope equipped with a Zeiss Axio Cam MRc 5 digital camera operated in differential interferential contrast (DIC) mode.

### Yolk granules flow cytometry

Suspensions of Ygs were obtained by gently disrupting freshly dissected chorionated oocytes in PBS (2 oocytes in 250 μl of buffer) using a plastic pestle. Population profiles of Ygs were acquired in a FACS Calibur equipment (BD Bioscience) powered by CellQuest Pro v5.1 software and analyzed using Flowing Software 2.5.1. For each biological replicate, samples were prepared using a pool of 2 eggs from different insects.

### Statistics

Student’s t-test was used for the comparison of two different conditions, and One-way ANOVA or Two-way ANOVA followed by Tukey’s Multiple Comparisons for the comparison among more than two conditions. Log-rank (Mantel-Cox) test was performed for the survival experiments. All the tests used the GraphPad Prism 8 software. Differences were considered significant at p < 0.05.

## Results

### Rhodnius prolixus harbors a conserved, single-copy VgR transcript that is highly enriched in early-stage oocytes

In *R. prolixus*, the sequence RPRC000551 was previously identified by [42] as the single isoform of the vitellogenin receptor (VgR) in this species. The VgR transcript comprises 5,454 nucleotides, encoding a putative protein of 1,817 amino acids. Domain analysis revealed that the predicted protein contains all conserved structural motifs typical of VgRs, including LDLa (Low-density lipoprotein receptor class A; pfam00057), EGF (Epidermal growth factor-like; pfam00008), LY (Low-density lipoprotein receptor YWTD; pfam00005), EGF_like (pfam07974), and EGF_CA (Calcium-binding EGF-like; pfam07645) domains, as well as a C-terminal transmembrane domain (Figure 1A). We next quantified vitellogenin receptor (VgR) transcript levels in different tissues of adult *R. prolixus* using RT-qPCR. In adult females, VgR expression was two orders of magnitude higher in the ovary compared with the levels detected in the midgut and fat body. Interestingly, in adult males, approximately 25% of the expression observed in the female ovary was detected in the testis, indicating that VgR transcription is not restricted to females. Moreover, the male fat body expressed about five times more VgR than the female fat body (Figure 1B). Within the ovary, VgR mRNA levels were 2–3 times higher in the tropharium and in pre-vitellogenic follicles than in vitellogenic and chorionated (mature) oocytes (Figure 1C). Together, these data indicate that VgR expression is predominantly female-and ovary-biased, but also present in male tissues, and dynamically regulated throughout oocyte development.

**Figure 1:**
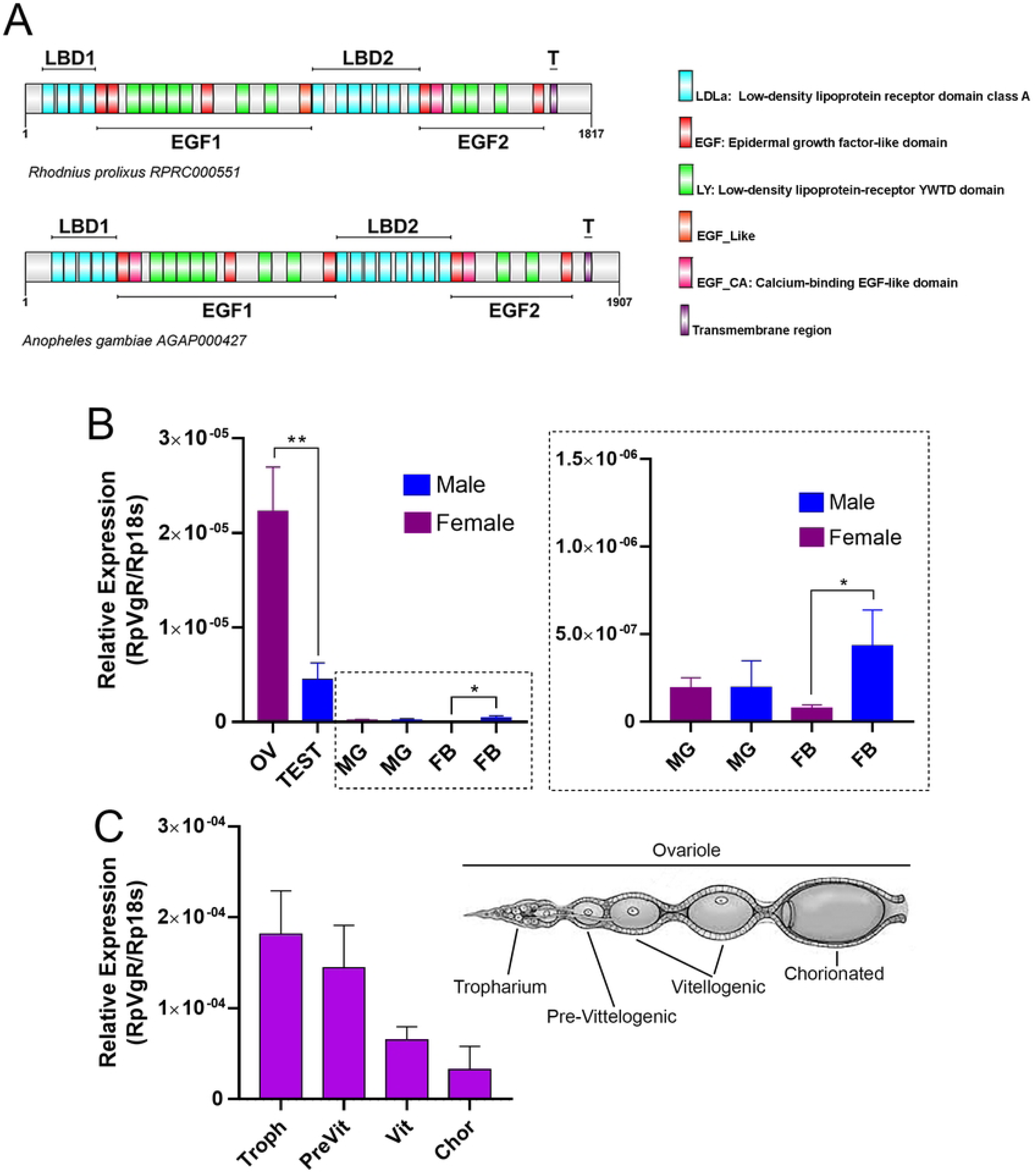
Rhodnius *prolixus* expresses a single conserved VgR isoform enriched in early oocytes. (A) Schematic diagram showing the conserved domain of *R. prolixus* and *Anopheles* SMART. (B) RT-qPCR showing the expression profile of VgR in the different organs of adult males and females dissected 7 days after blood feeding. VgR expression is significantly higher in the ovary (OV) and testis (TEST) when compared to the Fat body (FB) and midgut (MG) (n=6-10). (C) RT-qPCR showing the expression profile of VgR during oocyte development. Troph, tropharium; PreVit, pre-vitellogenic; Vit, vitellogenic; Chor, chorionated (n=3-4). A schematic representation of the ovariole is shown. Graphs show mean ± SEM. T-test, One-way ANOVA. *p<0.05, **p<0.01.

### Systemic VgR knockdown extends lifespan without disrupting vitellogenesis endocrine readouts

To further explore the role of VgR in the physiology of adult females, we synthesized and injected dsRNAs targeting VgR two days before blood feeding and monitored the resulting phenotypes throughout the subsequent gonotrophic cycle (Figure 2A). Injection of dsVgR achieved an efficient knockdown, with at least 80% reduction in VgR transcript levels across all analyzed organs (Figure 2B). We first assessed whether VgR silencing affected key aspects of vitellogenic physiology. Blood digestion and diuresis appeared unaffected, as no differences were observed in the rate of post-feeding weight loss between control and VgR-silenced females during the gonotrophic cycle (Figure 2C). Interestingly, a significant increase in longevity was detected: the median survival increased from 34.5 days in control females to 46 days in VgR-silenced insects (Figure 2D). Because juvenile hormone (JH) and ecdysone signaling are crucial for vitellogenesis [43–47], we next examined whether systemic VgR silencing altered the expression of these hormones’ response genes in the ovary. The JH-responsive gene Kr-h1 showed no significant changes following VgR knockdown (Figure 2E). In contrast, some ecdysone-responsive genes, including E74, BR-C, HR-3, and FTZ-F1, were upregulated by at least two-fold relative to controls, whereas E75 and HR-4 remained unaltered (Figures 2F–H). These results indicate that VgR silencing does not impair the activation of primary reproductive endocrine pathways in adult female *R. prolixus*.

**Figure 2.**
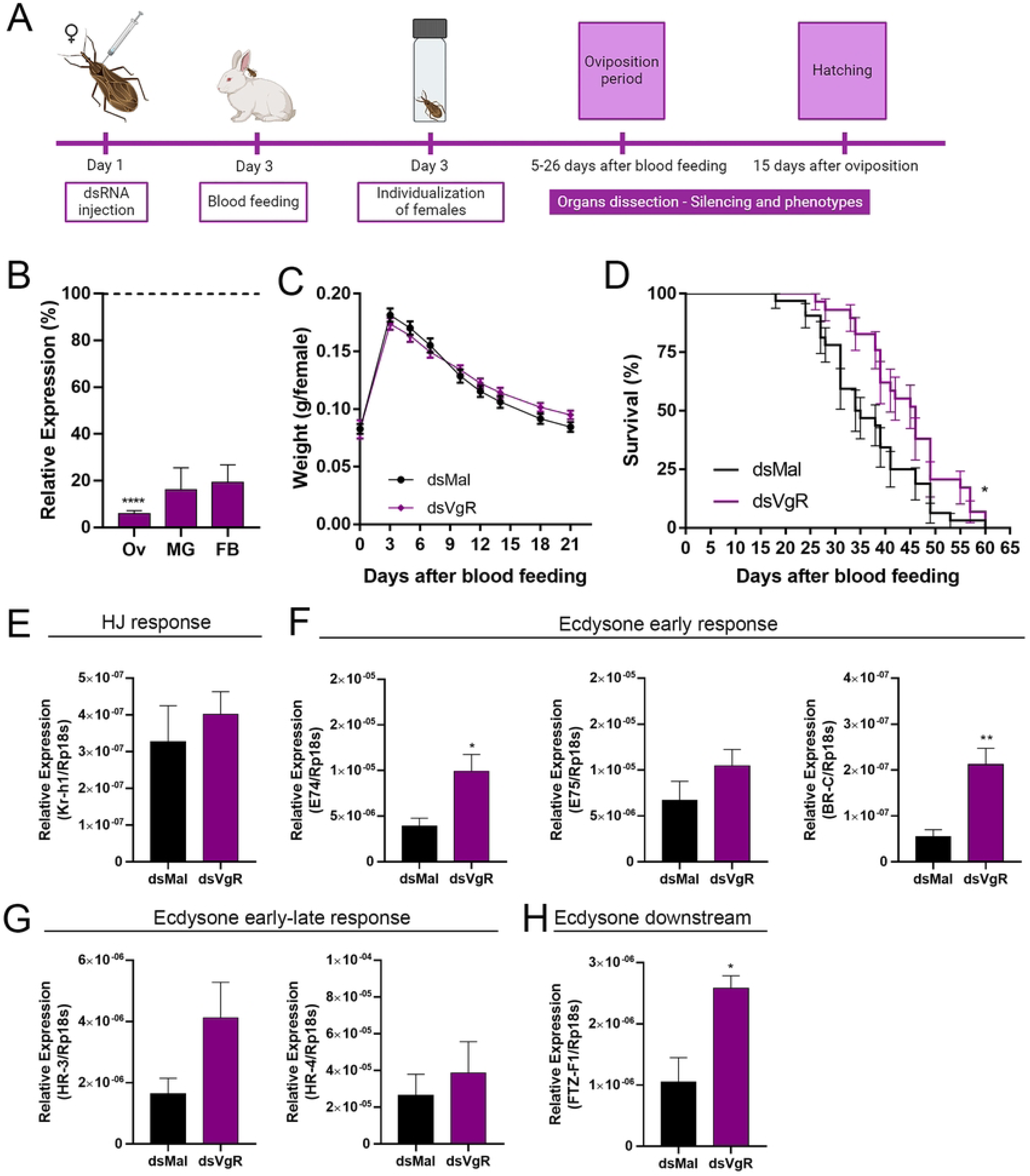
VgR knockdown extends female lifespan without impairing vitellogenic physiology. **(A)** Experimental design for dsRNA injection. Insects were injected 2 days before blood feeding, individualized, and the phenotypes were observed during the gonotrophic cycle. **(B)** Silencing efficiency 7 days after the blood meal is presented in different organs after injection of dsRNA designed to target VgR Fat body (FB), midgut (MG), and ovary (OV) (n=4-10). T-test. **(C)** Effects of VgR silencing on female digestion/diuresis, showing their weight during the gonotrophic cycle (n=30). Two-Way ANOVA. p>0.05. **(D)** Survival rates of control and VgR-silenced females (n=30). Log-rank (Mantel-COX) test. **(E)** RT-qPCR showing the expression profile of the HJ effector gene Kr-h1 in the ovary of control and VgR silenced females 7 days after blood feeding (n=5-6). T-test p>0.05. **(F-H)** RT-qPCR showing the expression profile of the ecdysone effector genes E74, E75, BR-C, HR3, HR-4 and FTZ-F1 in the ovaries of control and VgR silenced females 7 days after blood feeding (n=5-6). T-test. Graphs show mean ± SEM. *p<0.05, ****p<0.0001.

### VgR silencing blocks yolk uptake, causing hemolymph accumulation of Vg and RHBP

To further investigate the effects of VgR silencing, we quantified the total protein content in the hemolymph of vitellogenic females. VgR-silenced females contained approximately twice as much total protein as control insects (Figure 3A). Analysis of hemolymph protein composition by SDS-PAGE revealed a marked accumulation of vitellogenin (Vg) in VgR-silenced females. This accumulation was accompanied by an increase in Rhodnius heme-binding protein (RHBP), but not in the other major hemolymph proteins, lipophorin (Lp) [48,49] and arylphorin (Ar) [50] (Figure 3B). Consistent with these findings, the hemolymph of VgR-silenced [35,51–53]. The titration of total RHBP in the hemolymph was determined following the protocol described by [32]. Hemolymph samples were saturated by the addition of hemin, and absorbance spectra were recorded. The characteristic Soret peak at 412 nm confirmed the higher concentration of RHBP in dsVgR hemolymphs (Figure 3C). In dsMal hemolymphs, the titration profile indicated concentrations of 201.1 ± 89.9 µM RHBP. In dsVgR, the profile indicated a higher total amount of RHBP, reaching 403.1 ± 98.6 µM. Upon dissection, the ovaries of VgR-silenced females displayed an altered morphology, characterized by smaller, pale (white) oocytes, in contrast to the reddish, mature oocytes observed in control insects (Figures 3D–E). Together, these results indicate that VgR silencing does not affect Vg synthesis or secretion by the fat body but rather impairs the uptake of both Vg and RHBP by developing oocytes, demonstrating that RHBP internalization is functionally linked to Vg endocytosis, and consequently leads to their accumulation in the hemolymph.

**Figure 3:**
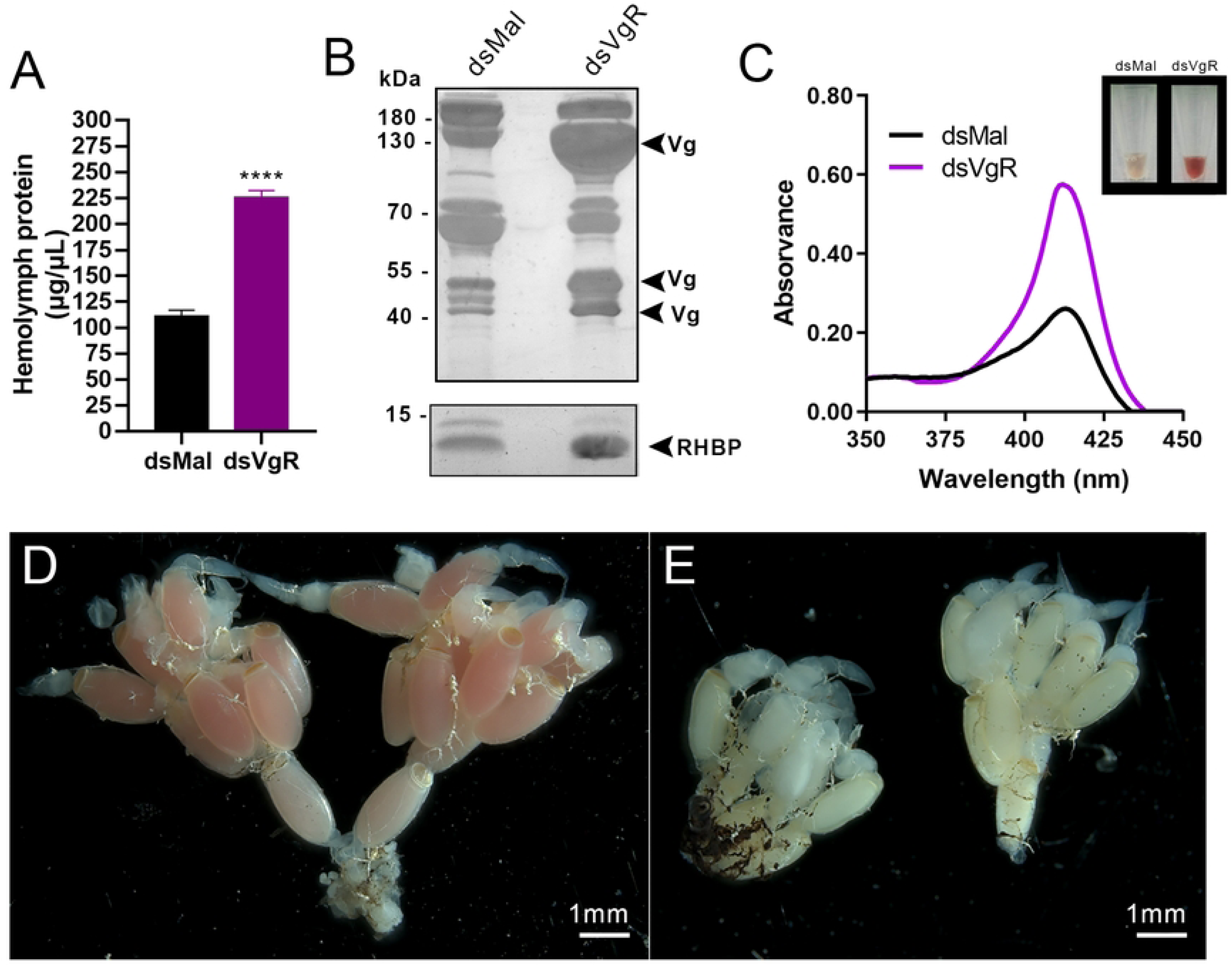
Impaired yolk uptake in VgR-silenced females leads to hemolymph accumulation of yolk proteins. **(A)** Control and silenced hemolymph protein quantifications 7 days after the blood meal (n=6-10). T-test. **(B)** 10% (upper panel) and 13,5% (lower panel) SDS-PAGEs of the hemolymphs of control and VgR-silenced females 7 days after the blood meal (n=6). Arrowheads point to the Vg subunits and RHBP. **(C)** Light absorption spectra of hemolymph from females injected with dsMal or dsVgR. RHBP Soret peak is shown at 412nm for both conditions. Inset shows a representative image of the hemolymphs collected on day 7 after blood feeding. The results are representative of two independent experiments. **(D-E)** Representative images of ovaries dissected from control and VgR silenced females 7 days after blood feeding. Scale bar: 1 mm. All graphs show mean ± SEM. ****p<0.0001.

### Oviposition rates persist after VgR knockdown, but yolk-depleted eggs fail to support embryogenesis

Despite the abnormal morphology of the oocytes, VgR-silenced females oviposited normally, with no significant differences in oviposition rates compared to control insects (Figure 4A). However, examination of the F1 eggs revealed severe phenotypic alterations: only 5% of the eggs laid by VgR-silenced females displayed normal morphology, while 95% were small and white (Figures 4B–C). Overall, these eggs exhibited low viability, with a total hatching rate of approximately 10% (Figure 4D). Among morphologically normal eggs, 79% of embryos were viable, whereas only 4% of embryos from the white eggs were successfully developed (Figure 4E). Despite their reduced size and lack of pigmentation, the white eggs were fertilized, as confirmed by PCR detection of a Y chromosome transcript, following the method previously described by [32](Figure 4F). Biochemical analyses revealed that eggs from VgR-silenced females exhibited an ∼80% reduction in total protein content, accompanied by drastically reduced levels of vitellin (Vt) and RHBP (Figure 4G). In contrast, despite the smaller egg volume, the total glycogen content of these eggs was only slightly reduced when compared to controls (Figure 4H). Lipid analysis by thin-layer chromatography (TLC) showed that triacylglycerol (TAG), hydrocarbon (HC), fatty acid (FA), and monoacylglycerol (MAG) showed similar levels in control and silenced groups (Figure 4I). However, the contents of diacylglycerol (DAG), cholesterol (CHO), and phospholipid (PL) were reduced by approximately 55%, 68%, and 45%, respectively (Figure 4I).

**Figure 4:**
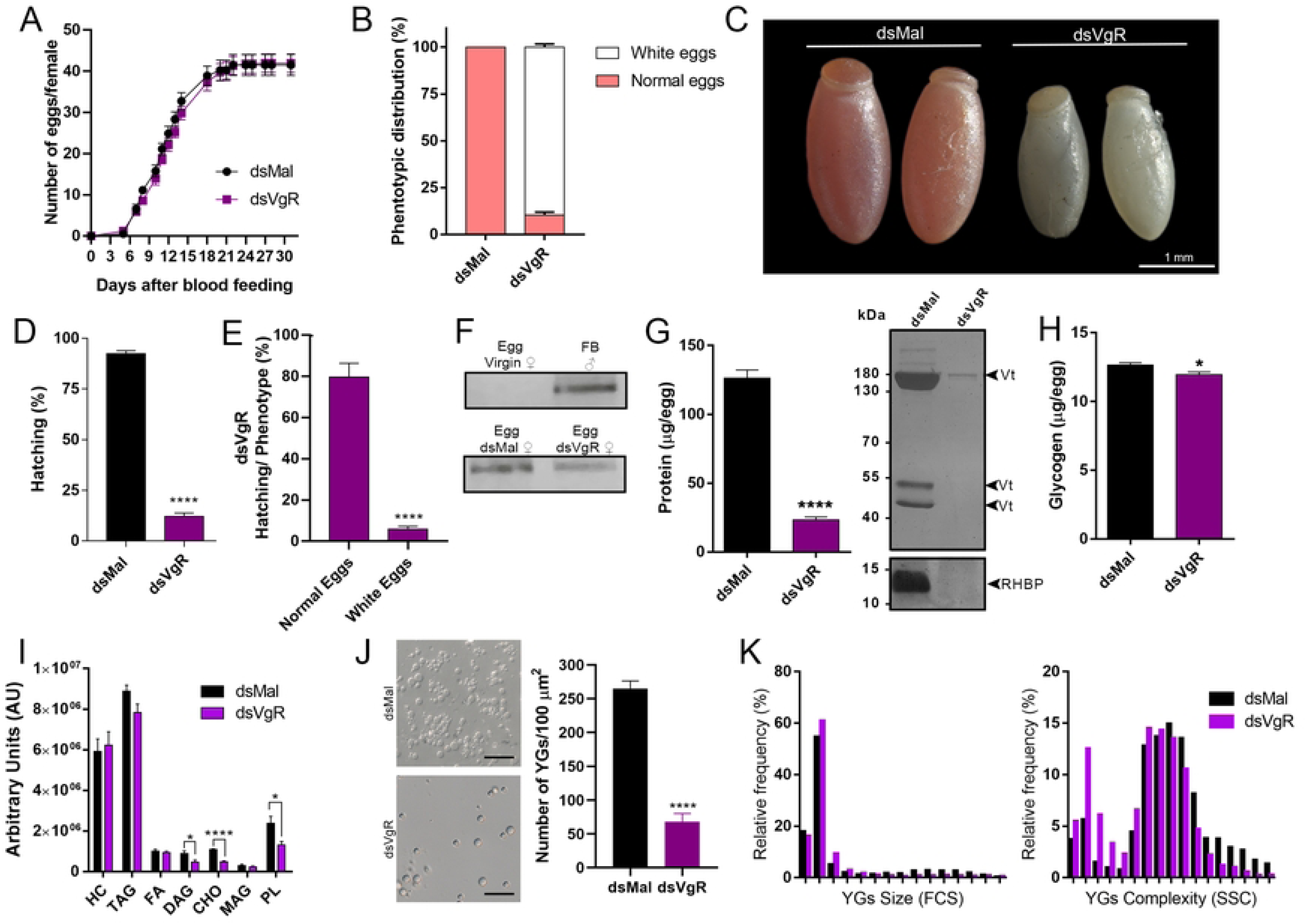
Yolk-depleted eggs from VgR-silenced females are fertilized but non-viable. **(A)** Oviposition rates control and silenced females during the gonotrophic cycle (n=30). Two-Way ANOVA p>0.05. **(B)** Phenotypic distribution of the eggs laid by control and VgR silenced insects (n=30). **(C)** Representative images of freshly laid eggs. Scale bar: 1 mm. **(D)** F1 total hatching rates after VgR silencing (n=30). T-test. **(E)** F1 hatching rates per phenotype (n=30). T-test. **(F)** Genomic DNA from eggs laid by control and VgR silenced females were extracted and used as templates for amplifying a specific Y chromosome fragment by PCR. Genomic DNA from nonfertilized eggs laid by virgin females was used as negative controls. Fat body DNA from adult male insects was used as a positive control. **(G)** Protein levels of freshly laid eggs from control and VgR-silenced females (n=4-7). T- test. SDS-PAGE protein profile of freshly laid eggs from control and VgR-silenced females (n=4-7). Arrowheads point to Vitellin (Vt) and RHBP. **(H)** Glycogen content quantified in eggs laid by control and VgR females (n=7). T-test. **(I)** Neutral lipid analysis was performed by TLC. Diacylglycerol (DAG), cholesterol (CHO), phospholipids (PL), Hydrocarbons (HC), triacylglycerol (TAG), monoacylglycerol (MAG) (n=8). T-test. **(J)** Representative images of the yolk granules (YGs) extracted from control and VgR-silenced females observed under the light microscope (n=2). Scale bar: 30 μm. Quantifications of the number of YGs extracted from dsMal and dsVgR oocytes (n=5). T-test. **(K)** The YGs suspensions were analyzed by flow cytometry. Frequency histogram analysis reveals a shift in the size (FSC) and complexity (SSC) of YGs in eggs laid by VgR-silenced females (n=16-22). All Graphs show mean ± SEM. *p<0.05, ****p<0.0001.

Together, these findings demonstrate that VgR silencing does not impair oviposition or fertilization, but severely disrupts yolk deposition, resulting in protein- and lipid-depleted eggs unable to sustain embryonic development. Because cholesterol (CHO) and phospholipids (PL) are major components of biological membranes, their reduction in VgR-silenced eggs, together with the observed yolk depletion, led us to hypothesize that yolk granule (YG) biogenesis was compromised. To test this, YG suspensions were prepared by directly disrupting the eggs on a glass slide and examining them under differential interference contrast (DIC) microscopy. We found that the number of YGs was reduced by approximately 70% in oocytes from VgR-silenced females compared to controls (Figures 4J). To further characterize the YG population, we used flow cytometry to analyze YGs isolated from chorionated oocytes of control and VgR-silenced females, as previously described by [54,55]. The analysis provided information on YG size (forward scatter, FSC) and internal complexity (side scatter, SSC). The resulting frequency histograms (Figures 4K) revealed that, although YGs from both groups displayed a wide range of sizes and complexities, those from VgR-silenced oocytes were predominantly smaller and less complex than those from controls. Together, these findings demonstrate that loss of VgR function impairs YG formation, resulting in defective YG biogenesis and supporting the idea that VgR-mediated endocytosis is essential for yolk accumulation during oocyte maturation.

### VgR silencing reshapes oocyte bacterial and core virome loads, and these effects are not explained by changes in Vg abundance in the oocyte or the hemolymph

Because yolk accumulation is known to participate in the shuttling of microbes to the oocyte and contributes to the vertical transmission of both pathogenic and non-pathogenic microbiota [26,27,56–60], we investigated bacterial and viral levels in yolk-depleted oocytes. In addition to *VgR* silencing, direct *Vg* silencing also produces yolk-depleted eggs that are morphologically and biochemically similar to those observed under *VgR* knockdown (Pereira et al., 2025). However, the physiological context differs markedly between the two conditions. As shown in Figure 5A, *VgR* silencing results in high circulating Vg, whereas *Vg* silencing leads to Vg-depleted hemolymph (Pereira et al., 2025). Because the hemolymph is likely the main route through which microbes reach the oocytes, we also asked whether bacterial and viral loads within the oocytes would reflect differences in Vg abundance in the hemolymph. To address this, we quantified bacterial and viral levels in both oocytes and hemolymph from *Vg*- and *VgR*-silenced insects (Figure 5). In oocytes from *VgR*-silenced insects, qPCR analysis of the bacterial 16S rRNA gene revealed a marked reduction in bacterial load (Figure 5B). Regarding the *R. prolixus* oocyte core virome [28], seven viruses were initially detected. However, RpV2 levels were near the detection threshold and were therefore excluded from further analyses. Among the six remaining viruses, *VgR*-silenced oocytes displayed increased levels of RpV4, RpV5, RpV6, and RpV7 (Figure 5D–H). In contrast, the reduction in bacterial load was less pronounced in *Vg*-silenced eggs compared with those lacking *VgR* (Figure 5B). For the viruses, all members except RpV7 tended to decrease in *Vg*-silenced oocytes (Figure 5D–H).

**Figure 5.**
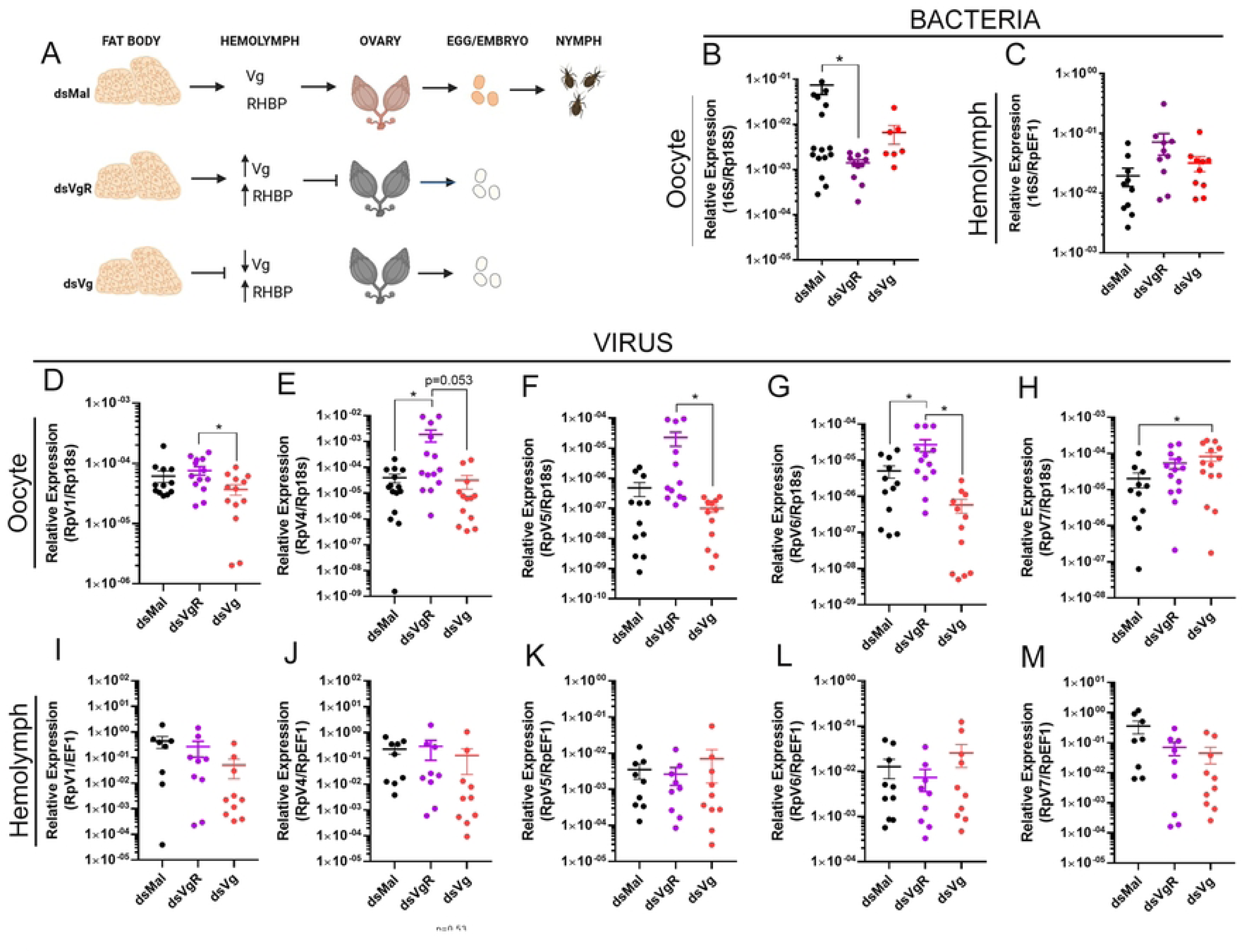
VgR knockdown reshapes bacterial and viral composition of oocytes. (A) Schematic representation of hemolymph, ovary, and embryo phenotypes following *VgR* or *Vg* silencing in *R. prolixus*. Vg data are from Pereira et al., 2025. *VgR* silencing leads to accumulation of Vg in the hemolymph, yolk-depleted eggs, and unviable embryos. *Vg* silencing results in Vg-depleted hemolymph, yolk-depleted eggs, and unviable embryos. In both cases, RHBP levels are increased in the hemolymph. (B) RT-qPCR analysis of bacterial 16S rRNA levels in oocytes from *VgR*- and *Vg*-silenced females (n=7-16). (C) RT-qPCR analysis of bacterial 16S rRNA levels in the hemolymph (n=10). (D–H) RT-qPCR quantification of different viral components of the core oocyte virome of R. prolixus in oocytes from *VgR-* and *Vg*-silenced females (n=12-13). (I–M) RT-qPCR quantification of the same viral components in the hemolymph (n=10). All graphs show mean ± SEM. One-way ANOVA; *p < 0.05.

Since both *Vg-* and *VgR-*silencing result in morphologically and biochemically similar yolk-depleted eggs, we hypothesized that the observed differences in bacterial and viral levels between Vg- and VgR-silenced females might reflect alterations occurring in the hemolymph. To test this possibility, we quantified bacterial and viral loads in the hemolymphs of both conditions. Hemolymph samples exhibited more dispersed bacterial (Figure 5C) and viral levels (Figure 5 I-M), without consistent trends correlating with those observed in the oocytes.

### Bacterial and viral load changes are not explained by direct Vg antimicrobial activity

Previous studies have shown that purified Vg from species such as *Apis mellifera*, *P. longicornis*, *C. carpio*, and *B. japonicum* were experimentally tested against bacterial strains, typically *E. coli* and/or *S. aureus*, showing direct antibacterial activity (e.g., growth inhibition, membrane disruption, or bacteriostatic effects) [60]. To test whether *R. prolixus* Vg displays antibacterial activity, purified Vg was incubated with *E. coli*, and *S. aureus* cultures for 24 h at 37°C. Bacterial growth was monitored and relative growth was calculated against untreated controls. Across all concentrations tested (12.5–200 µg/ml), *E. coli* growth remained between 91–99% of control levels (Table S2), while *S. aureus* growth ranged from 99–108% of controls (Table S3). No concentration-dependent inhibition was observed for either bacterial species, indicating that purified *R. prolixus* Vg does not exert detectable bacteriostatic or bactericidal effects under these conditions.

Viral infectivity was evaluated using a standard plaque assay for Mayaro virus (MAYV), and the number of plaque-forming units (PFU/mL) was quantified to determine viral titters. No significant differences in plaque number or morphology were observed between control and Vg-treated (400 µg/ml) groups, with similar PFUs for both conditions, indicating that Vg treatment did not alter MAYV infectivity or replication efficiency under the tested conditions (Figure S1). Altogether, the data indicates that the observed alterations in viral and bacterial loads are unlikely to result from the direct action of Vg within the oocytes.

### Ovaries are immunocompetent, but defensin and RNAi pathway modulation do not explain microbial shifts in VgR knockdown oocytes

Another possibility we considered to explain the observed microbial changes was that *VgR* silencing might influence the insect’s immune response, either systemically or within the immunocompetent. Under pathogen-unchallenged conditions, qPCR revealed that the expression of *Defensin A* and *Defensin B* was approximately 25% and 80% higher, respectively, in the ovary than in the fat body, whereas *Prolixicin* exhibited a basal expression roughly tenfold greater in the fat body compared to the ovary (Figure 6A). *Defensin C* was not detected under these conditions (data not shown). To further evaluate immune activation, we analyzed the response of these antimicrobial peptides to infections with *Trypanosoma rangeli*, using a protocol previously described by [37]. Both the fat body and the ovary upregulated all three *defensin* genes upon infection, reaching comparable induction levels in both tissues (Figure 6B). These results demonstrate that, in addition to the fat body, the ovary of *R. prolixus* is an immune-responsive tissue capable of mounting a transcriptional response to immune challenge.

**Figure 6.**
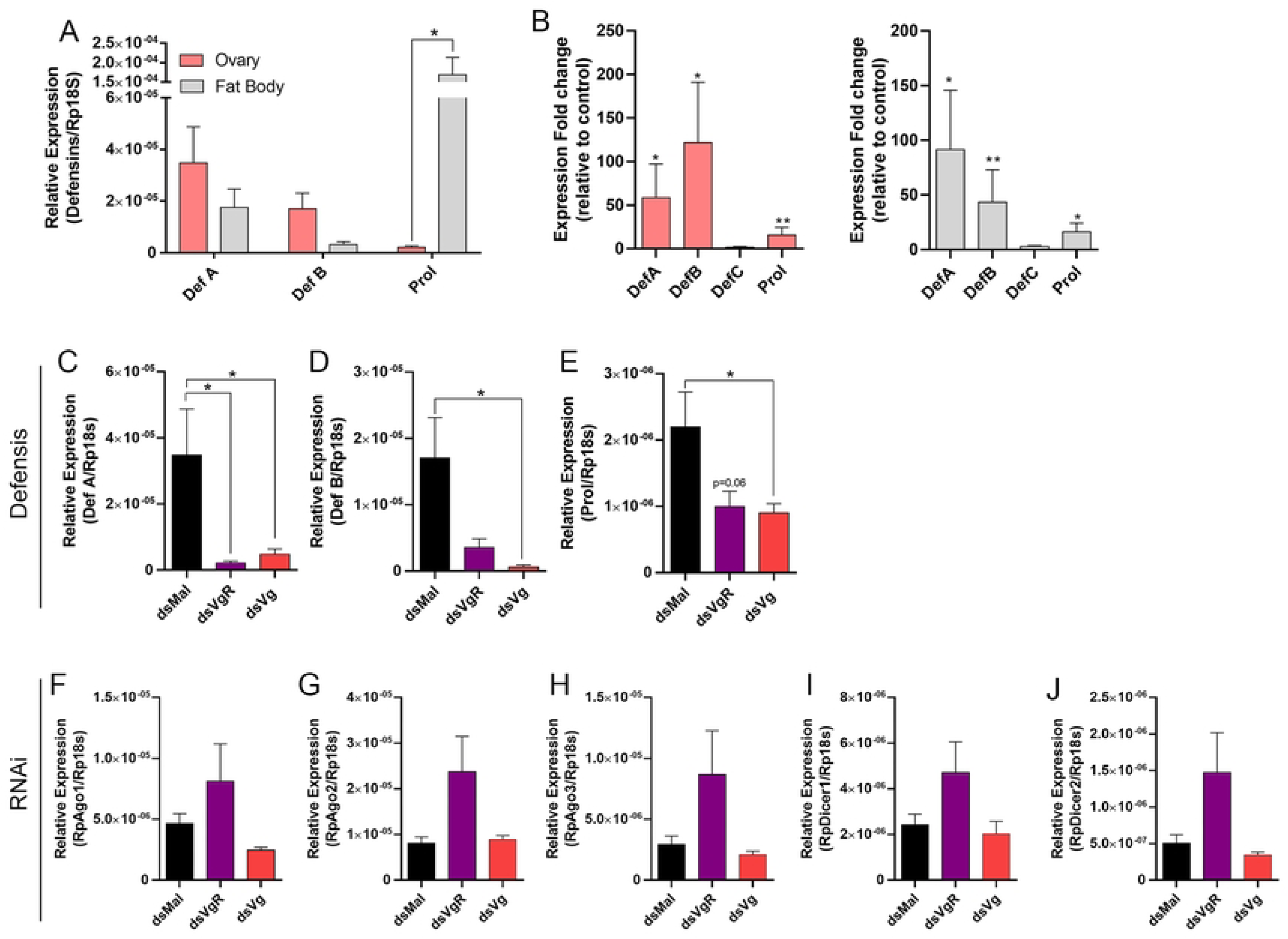
Ovary immune responses are present, but defensin and RNAi modulation do not explain microbial shifts. **(A)** RT-qPCR showing the basal expression levels of *Defensin A*, *Defensin B*, and *Prolixicin* in the fat body and ovary of *R. prolixus* (n=8-9). **(B)** Induction of the same antimicrobial peptide genes in the fat body and ovary after *Trypanosoma rangeli* infection (n=6). **(C–E)** Expression of *Defensin A*, *Defensin B*, and *Prolixicin* in the ovaries of *VgR*- and *Vg*-silenced females (n=8-9). (F–J) Expression of core RNA interference pathway components (*Dicer-1*, *Dicer-2*, *Argonaute-1*, *Argonaute-2*, and *Argonaute-3*) in the ovaries of *VgR*- and *Vg*-silenced females (n=8-9). All graphs show mean ± SEM. One-way ANOVA; *p < 0.05.

Having established that the ovary is immunocompetent, we next examined *defensin* expression in *VgR*- and *Vg*-silenced females to determine whether the observed changes in bacterial loads were associated with altered immune response. In both knockdowns, the expression levels of all *defensin* genes were markedly reduced in the ovary (Figure 6 C-E), a pattern that correlated negatively with the decreased bacterial load detected by 16S rRNA qPCR. Thus, modulation of canonical immune effectors is unlikely to account for the differences in bacterial abundance observed in the oocytes of *VgR*- and *Vg*-silenced insects.

Brito et al. (2021) [28] demonstrated that RNA interference (RNAi) represents the primary mechanism controlling the expansion of the *R. prolixus* core virome during oogenesis. In that study, the authors detected abundant 22-nucleotide viral small interfering RNAs (vsiRNAs) targeting all members of the core virome and proposed that vsiRNA-mediated silencing is likely the main mechanism regulating viral load in the oocytes. In this system, viral double-stranded RNA intermediates are processed by Dicer into vsiRNAs, which are loaded into Argonaute-containing RISC complexes, where Argonaute acts as the catalytic slicer that cleaves complementary viral RNAs to suppress replication. To further investigate the basis of the viral load alterations observed in our knockdown oocytes, we examined the expression of key RNAi components to determine whether this pathway might be modulated in *VgR*- and *Vg*-silenced females. Interestingly, *VgR* silencing, but not *Vg* silencing, was associated with a tendency toward upregulation of all *Argonaute* and *Dicer* isoforms (Figure 6 F-J), concomitant with the increased viral loads detected in these oocytes. Thus, a modulation in the RNAi pathway does not appear to be the cause of the observed differences in viral abundance, suggesting that other regulatory mechanisms may influence virome dynamics in *VgR*- and *Vg*-silenced ovaries.

### VgR is not transcriptionally modulated in Vg-silenced oocytes

Next, we hypothesized that the opposite patterns of viral abundance observed between *VgR*- and *Vg*-silenced oocytes could be related to the differential presence of *VgR*. Specifically, we reasoned that oocytes from *Vg*-silenced females might up-regulate *VgR* expression to compensate for the reduced availability of circulating Vg. In this scenario, the lower viral loads detected in *Vg*-silenced oocytes would result from increased *VgR* expression. To test this hypothesis, we quantified *VgR* transcript levels in *Vg*-silenced oocytes and found them to be similar to control samples (Figure 7), indicating that *VgR* expression is not up-regulated under Vg-deficient conditions and is likely not associated with the Vg-specific changes observed in bacterial and viral loads.

**Figure 7.**
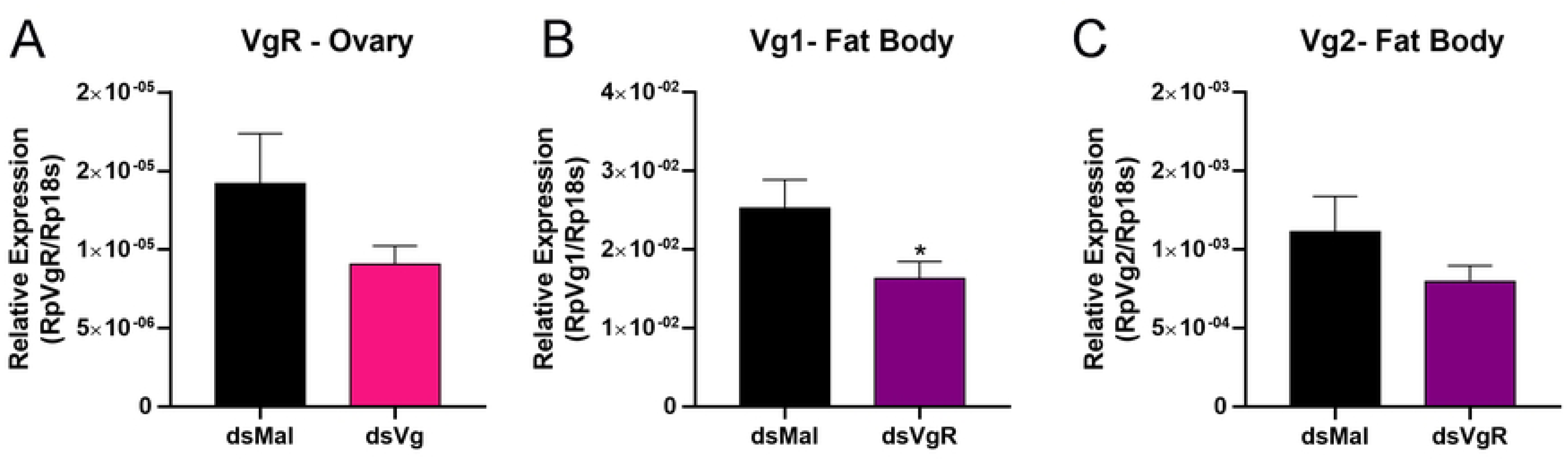
VgR is not transcriptionally modulated in Vg-silenced oocytes. (A) RT-qPCR quantification of *VgR* transcript levels in the ovaries of *Vg*-silenced females (n=8-9). (B-C) RT-qPCR quantification of Vg1 and Vg2 transcript levels in the fat body of *VgR*-silenced females (n=8-9). Graphs show mean ± SEM. T-test; *p < 0.05.

### Male VgR is dispensable for fertility and progeny viability

Although VgR is classically associated with female reproduction, evidence from other species indicates that it may also be expressed in males [22,61–63]. Consistent with this, we detected VgR expression in adult male tissues, with the testis exhibiting approximately 20% of the expression levels observed in the female ovary (Figure 8A). To explore potential male-related two days before blood feeding, following the same protocol used for females. Efficient knockdown was achieved, with at least 65% reduction in VgR transcript levels in the fat body, midgut, and testis (Figure 8B). Despite this, VgR silencing did not affect blood digestion over 21 days (Figure 8C) or male longevity (Figure 8D).

**Figure 8:**
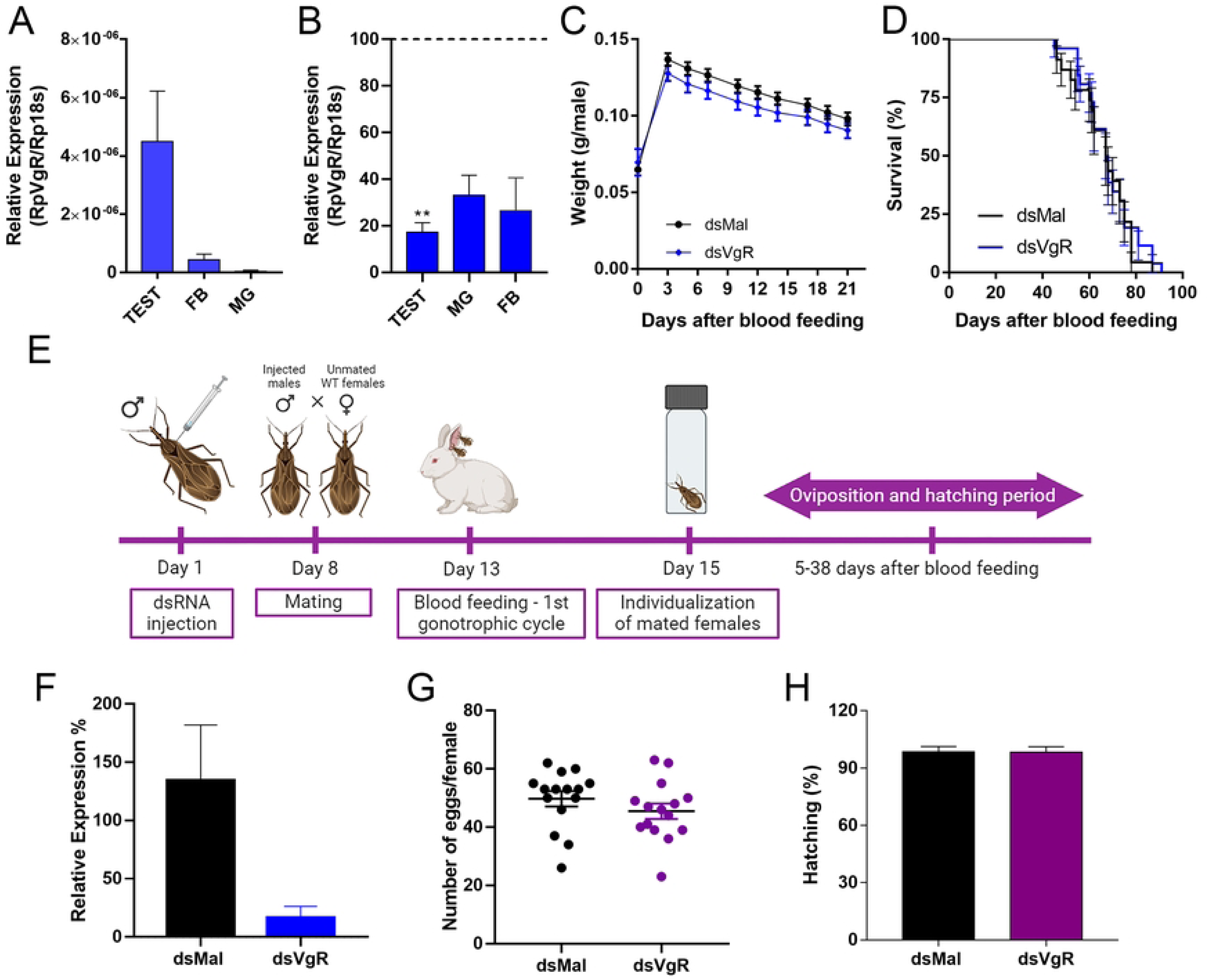
Male VgR expression is dispensable for fertility and progeny viability. **(A)** RT-qPCR showing the expression profile of VgR in the different tissues of adult males dissected 7 days after blood feeding (n=5-8). One-way ANOVA. **(B)** RT-qPCR shows the silencing efficiency for VgR in different organs after injection of dsRNA designed to target VgR (n=3-7). t-test. Fat body (FB), midgut (MG), and testis (TEST). **(C)** Effects of VgR silencing on male digestion/diuresis, showing their weight during the gonotrophic cycle (n=30). Two-Way ANOVA. p>0.05. **(D)** Survival rates of control and Vg-silenced females (n=30). Log-rank (Mantel-COX) test. p>0.05. **(E)** Experimental design for dsRNA injections in the mating experiments. Adult males were injected with VgR dsRNA 13 days before blood feeding. On day 8 after dsRNA injection, the silenced males were allowed to mate with virgin females for 5 days. After the mating period, both males and females were blood-fed. **(F)** Silencing efficiency for the males on day 7 after dsRNA injection (before mating) is presented in different organs (n=5). t-test. p>0.05. Testis (TEST). **(G)** Total oviposition of females mated with control and VgR silenced males (n=15). T-test p>0.05. **(H)** Hatching rates of females mated with control and VgR silenced males (n=15). *t*-test. p>0.05. All graphs show mean ± SEM.*p<0.05, **p<0.01.

Given that Vg silencing in male *Chrysopa pallens* has been reported to reduce oviposition and hatching rates in their mated females [64], we tested whether a similar phenomenon occurs in *R. prolixus* when silenced for VgRs. To this end, we adapted a mating protocol in which VgR-silenced males were paired with wild-type virgin females seven days post-injection and allowed to mate for five days before blood feeding (Figure 8E). Knockdown efficiency was confirmed by qPCR, showing at least 85% VgR silencing in the testis (Figure 8F). Despite the silencing efficiency, female oviposition rates following mating with VgR-silenced males remained unchanged (Figure 8G). Similarly, the hatching rates of the laid eggs were unaffected (Figure 8H). Together, these results indicate that although VgR is expressed at lower levels in male *R. prolixus*, its silencing does not impact male physiology, fertility, or offspring development, suggesting that male VgR plays no essential reproductive role in this species.

## Discussion

Vector-borne NTDs remain a major global health challenge, with Chagas disease affecting millions across Latin America and emerging globally [65,66]. *Rhodnius prolixus*, a key vector and a classical model in insect physiology [3], offers some experimental advantages, including a fully sequenced genome [2] and extensive transcriptomic resources for digestive and reproductive tissues [29,67–70]. Its robust RNAi response further supports its use in dissecting reproductive pathways and identifying molecular targets to reduce vector fertility.

Our findings show that *R. prolixus* expresses a single, structurally conserved VgR that is predominantly localized to the ovary, consistent with previous descriptions in other insects [7]. The high expression of VgR in early oocytes supports its role in receptor-mediated yolk uptake during vitellogenesis. Interestingly, VgR transcripts were also detected in the testis, suggesting that this receptor may have additional, yet unexplored, functions in male physiology, as also observed in other insect species [22,61–63].

Despite the clear reproductive association, the phenotypic consequences of VgR loss in females revealed an unexpected dissociation between yolk accumulation and oviposition rates. Indeed, despite the dramatic impact of VgR silencing on yolk uptake, oviposition rates and fertilization were not affected, revealing that the completion of oogenesis is not contingent upon the actual accumulation of yolk. This indicates that the endocrine cues that define oocyte differentiation act independently of yolk load. This observation is consistent with results from Vg-silenced females that also lay the same number of yolk-depleted eggs [71], and refines our understanding of the mechanisms that control oogenesis in triatomines.

Although VgR silencing caused a marked depletion of yolk proteins, the other major energy reserves for the embryo, glycogen and TAG, remained unchanged, indicating that their accumulation occur independently of the VgR pathway. Yet, these reserves alone are apparently insufficient to sustain embryonic development. Notably, phospholipid and cholesterol levels, essential for membrane formation, were reduced in eggs from VgR-silenced females, likely because of the impaired buildup of internal membranes to sustain the YG biogenesis. Plus, DAG and phospholipids are known constituents of Vg [5], so the lack of Vg accumulation might also account for some of the observed reduction of those lipids.

The loss of VgR also led to alterations in the microbial profile of the oocytes. The reduction in bacterial 16S rRNA levels and the concomitant expansion of several members of the core virome highlight a potential link between yolk endocytosis and the vertical transmission of microbial components. In several insects, both bacteria and viruses have been shown to exploit reproductive pathways to reach developing oocytes. For instance, in *Drosophila melanogaster*, the endogenous retrovirus *Zam* is translocated from the follicular epithelium to the oocyte via vitellogenin trafficking [72,73]. Similarly, in the green rice leafhopper *Nephotettix cincticeps*, the rice dwarf virus attaches to the endosymbiont *Sulcia* to facilitate oocyte entry [74], while in *Laodelphax striatellus*, the rice stripe virus hijacks the VgR pathway to achieve vertical transmission [26,27]. In whiteflies, the tomato yellow leaf curl virus binds directly to vitellogenin to access the ovaries [75,76]. Vitellogenin itself is known to bind bacteria via pathogen-associated molecular patterns (PAMPs), functioning as a carrier molecule that can mediate bacterial transport into the oocyte [59,77].

Interestingly, we found that VgR silencing favored viral expansion while decreasing bacterial abundance in the oocytes, a pattern not recapitulated by *Vg* silencing. Instead, *Vg*-silenced oocytes exhibited reduced viral loads and only a mild decrease in bacterial abundance. Because both knockdowns generate yolk-depleted eggs, the distinct microbial patterns cannot be explained by the absence of yolk proteins in the oocytes per se. Differences in circulating Vg concentrations between the two conditions also failed to account for the observed phenotypes, as viral and bacterial loads in the hemolymph were comparable. Furthermore, purified *R. prolixus* Vg showed no direct antimicrobial or antiviral activity against *E. coli*, *S. aureus*, or Mayaro virus *in vitro*. In addition, neither immune gene modulation (defensins) nor RNAi pathway activation appears to be the primary cause of the microbial changes, as both correlated negatively with bacterial and viral loads. These negative correlations instead suggest that their modulation reflects a response to shifts in microbial levels rather than the driver of those changes. Altogether, these observations indicate that the altered microbial balance arises from the absence of functional VgR itself, rather than from yolk depletion or modulation of canonical immune pathways.

Assuming that the main determinants of the microbial alterations observed are VgR-related, it remains challenging to discern whether these changes arise primarily from altered entry (flux) of the microbes into the oocytes, or whether the differential flux of other components secondarily results in altered microbial expansion within the oocyte. Nevertheless, some mechanistic possibilities can be proposed. One possibility is that VgR itself acts as an entry or regulatory platform for additional yet unidentified, ligands that influence microbial flux and/or the maintenance within the oocyte. In this scenario, the absence of VgR would prevent the entry of one or more factors that normally disfavor viral expansion while sustaining bacterial presence, thereby shifting the microbial balance toward reduced bacterial load and enhanced viral abundance. Another possibility is that the reduction in VgR likely causes a severe alteration of the endocytic environment of the oocyte cortex, since the loss of endocytic activity required for the massive YG biogenesis probably disrupts the balance between membrane influx and recycling of surface proteins. Consequently, the repertoire of surface proteins available to mediate uptake events would change, potentially resulting in distinct fluxes of ligands or microbial particles into the oocyte through alternative, non-VgR-mediated routes.

Plus, when comparing VgR- and Vg-silencing data, the opposite viral trends observed in Vg-silenced oocytes when compared to VgR-silenced oocytes may also reflect differences in VgR localization and/or availability at the oocyte membrane. Under Vg silencing, the lack of VgR’s major ligand (Vg) likely increases the number of unoccupied VgR molecules at the oocyte surface, which could enhance receptor activity and facilitate the internalization of other VgR-interacting molecules, thus resulting in the opposite effect of silencing VgR. Moreover, the reduction in endocytic flux may further increase the residence time of VgR at the membrane. In this context, the higher abundance of unoccupied VgR molecules in the oocyte surface could underlie the opposing viral outcomes between VgR and Vg knockdowns, suggesting that the changes observed upon Vg silencing might stem from increased VgR availability to alternative ligands. Altogether, these observations suggest that VgR functions as a central node connecting endocytic dynamics with microbial accumulation in the developing oocyte, acting not only as a cargo carrier but also as a signaling receptor that coordinates these processes.

Although it remains challenging to design experimental approaches capable of disentangling the factors that define these microbial changes, whether they stem from altered entry, expansion, or survival within the oocyte. It would also be particularly interesting to explore how such alterations may influence vertical transmission and progeny viability. Brito et al. (2021) observed that all RpVs identified were markedly reduced during *R. prolixus* embryogenesis and proposed that this decline results from an upregulated RNAi response aimed at lowering viral levels to support normal embryo development, while still maintaining residual viral populations that later re-expand during the early nymphal stages. If the balance of viral load is indeed finely tuned to sustain potential symbiotic benefits, then the crosstalk between yolk accumulation and microbial inheritance may represent a key mechanism not only for ensuring progeny fitness but also for shaping broader aspects of species ecology, including habitat adaptation, interspecific interactions, and long-term vector-microbe coevolution.

Together, these findings suggest that VgR function extends beyond yolk accumulation to encompass a broader role in the molecular crosstalk between reproduction and microbial inheritance. This expanded functional framework highlights an unrecognized layer of complexity in the reproductive biology of triatomines.

## Acknowledgments

The authors thank Bruna Beatris Santana Afonso, Maysa Moura Lopes and Geane Cleia Pereira Braz for insect handling and care and CENABIO-UFRJ for providing microscopy equipment and facilities.

---

**Figure S1: Purified Vg does not affect Mayaro virus infectivity in cell culture. (A)** Representative plaque assay showing *Mayaro virus* (MAYV) infection in Vero cell monolayers incubated with vehicle (Control) or purified Vg (400 µg/ml). Viral plaques were visualized by crystal violet staining. **(B)** Quantification of viral titers expressed as plaque-forming units per milliliter (PFU/mL). Bars represent mean ± SEM from three independent experiments (n=3). t-test; *p* > 0.05.

## References

1. WHO. Vector-borne diseases. 2 Mar 2020 [cited 10 Jun 2022]. Available: https://www.who.int/news-room/fact-sheets/detail/vector-borne-diseases

2. Mesquita RD, Vionette-Amaral RJ, Lowenberger C, Rivera-Pomar R, Monteiro FA, Minx P, et al. Genome of Rhodnius prolixus, an insect vector of Chagas disease, reveals unique adaptations to hematophagy and parasite infection. Proceedings of the National Academy of Sciences. 2015;112: 14936–14941.

3. Lange AB, Leyria J, Orchard I. The hormonal and neural control of egg production in the historically important model insect, Rhodnius prolixus: A review, with new insights in this post-genomic era. Gen Comp Endocrinol. 2022; 114030.

4. Tufail M, Raikhel AS, Takeda M. Biosynthesis and processing of insect vitellogenins. Progress in vitellogenesis Reproductive biology of invertebrates. 2005;12: 1–32.

5. Raikhel AS, Dhadialla TS. Accumulation of yolk proteins in insect oocytes. Annu Rev Entomol. 1992;37: 217–251. doi:10.1146/annurev.en.37.010192.001245

6. Tufail M, Takeda M. Molecular characteristics of insect vitellogenins. J Insect Physiol. 2008;54: 1447–1458.

7. Tufail M, Takeda M. Insect vitellogenin/lipophorin receptors: molecular structures, role in oogenesis, and regulatory mechanisms. J Insect Physiol. 2009;55: 88–104.

8. Snigirevskaya ES, Raikhel AS. Receptor-mediated endocytosis of yolk proteins in insect oocytes. Progress in vitellogenesis Reproductive biology of invertebrates. 2005;12: 199–228.

9. Ramos I, Machado E, Masuda H, Gomes F. Open questions on the functional biology of the yolk granules during embryo development. 2022; 1–9. doi:10.1002/mrd.23555

10. Tufail M, Takeda M. Molecular cloning and developmental expression pattern of the vitellogenin receptor from the cockroach, Leucophaea maderae. Insect Biochem Mol Biol. 2007;37: 235–245.

11. Tufail M, Takeda M. Molecular cloning, characterization and regulation of the cockroach vitellogenin receptor during oogenesis. Insect Mol Biol. 2005;14: 389–401.

12. Cho K, Raikhel AS. Organization and developmental expression of the mosquito vitellogenin receptor gene. Insect Mol Biol. 2001;10: 465–474.

13. Schonbaum CP, Perrino JJ, Mahowald AP. Regulation of the vitellogenin receptor during Drosophila melanogaster oogenesis. Mol Biol Cell. 2000;11: 511–521.

14. Wang C, Bao H, Yan Z, Wang J, Wang S, Li Y. Knockdown of vitellogenin receptor based on minute insect RNA interference methods affects the initial mature egg load in the pest natural enemy Trichogramma dendrolimi. Insect Sci. 2025;32: 487–500.

15. Nobre IC da S, Coelho RR, de Souza FMC, Reis MA, Torres JB, Antonino JD. Insights from different reproductive gene knockdowns via RNA interference in the lady beetle Eriopis connexa: Establishing a new model for molecular studies on natural enemies. Arch Insect Biochem Physiol. 2024;116: e22125.

16. Nian X, Luo Y, He X, Wu S, Li J, Wang D, et al. Infection with ‘Candidatus Liberibacter asiaticus’ improves the fecundity of Diaphorina citri aiding its proliferation: a win-win strategy. Mol Ecol. 2024;33: e17214.

17. Lian Y, Peng S, Jia J, Li J, Wang A, Yang S, et al. Function of Vitellogenin receptor gene in reproductive regulation of Zeugodacus cucurbitae (Coquillett) after short-term high-temperature treatment. Front Physiol. 2022;13: 995004.

18. Han S, Wang D, Song P, Zhang S, He Y. Molecular characterization of vitellogenin and its receptor in Spodoptera frugiperda (JE Smith, 1797), and their function in reproduction of female. Int J Mol Sci. 2022;23: 11972.

19. Han H, Han S, Qin Q, Chen J, Wang D, He Y. Molecular identification and functional characterization of vitellogenin receptor from Harmonia axyridis (Coleoptera: Coccinellidae). J Econ Entomol. 2022;115: 325–333.

20. Jing Y-P, Wen X, Li L, Zhang S, Zhang C, Zhou S. The vitellogenin receptor functionality of the migratory locust depends on its phosphorylation by juvenile hormone. Proceedings of the National Academy of Sciences. 2021;118: e2106908118.

21. Ciudad L, Piulachs M, Bellés X. Systemic RNAi of the cockroach vitellogenin receptor results in a phenotype similar to that of the Drosophila yolkless mutant. FEBS J. 2006;273: 325–335.

22. Sheng Y, Chen J, Jiang H, Lu Y, Dong Z, Pang L, et al. The vitellogenin receptor gene contributes to mating and host-searching behaviors in parasitoid wasps. iScience. 2023;26.

23. Kaltenpoth M, Flórez L V., Vigneron A, Dirksen P, Engl T. Origin and function of beneficial bacterial symbioses in insects. Nature Reviews Microbiology. Nature Research; 2025. pp. 551–567. doi:10.1038/s41579-025-01164-z

24. Rodríguez J, Pavía P, Montilla M, Puerta CJ. Identifying triatomine symbiont Rhodococcus rhodnii as intestinal bacteria from Rhodnius ecuadoriensis (Hemiptera: Reduviidae) laboratory insects. Int J Trop Insect Sci. 2011;31: 34–37. doi:10.1017/S1742758411000014

25. Longdon B, Jiggins FM. Vertically transmitted viral endosymbionts of insects: Do sigma viruses walk alone? Proceedings of the Royal Society B: Biological Sciences. Royal Society; 2012. pp. 3889–3898. doi:10.1098/rspb.2012.1208

26. He K, Lin K, Ding S, Wang G, Li F. The vitellogenin receptor has an essential role in vertical transmission of rice stripe virus during oogenesis in the small brown plant hopper. Pest Manag Sci. 2019;75: 1370–1382. doi:10.1002/ps.5256

27. Huo Y, Yu Y, Liu Q, Liu D, Zhang M, Liang J, et al. Rice stripe virus hitchhikes the vector insect vitellogenin ligand-receptor pathway for ovary entry. Philosophical Transactions of the Royal Society B: Biological Sciences. 2019;374. doi:10.1098/rstb.2018.0312

28. De Brito TF, Coelho VL, Cardoso MA, De Abreu Brito IA, Berni MA, Zenk FL, et al. Transovarial transmission of a core virome in the Chagas disease vector Rhodnius prolixus. PLoS Pathog. 2021;17. doi:10.1371/journal.ppat.1009780

29. Leyria J, Orchard I, Lange AB. What happens after a blood meal? A transcriptome analysis of the main tissues involved in egg production in Rhodnius prolixus, an insect vector of Chagas disease. PLoS Negl Trop Dis. 2020;14: e0008516.

30. Livak KJ, Schmittgen TD. Analysis of relative gene expression data using real-time quantitative PCR and the 2(-Delta Delta C(T)) Method. Methods. 2001;25: 402–408. Available: http://www.ncbi.nlm.nih.gov/entrez/query.fcgi?cmd=Retrieve&db=PubMed&dopt=Citation&list_uids=11846609

31. Bustin SA, Benes V, Garson JA, Hellemans J, Huggett J, Kubista M, et al. The MIQE guidelines: Minimum information for publication of quantitative real-time PCR experiments. Clin Chem. 2009;55: 611–622. doi:10.1373/clinchem.2008.112797

32. Walter-Nuno AB, Oliveira MP, Oliveira MF, Gonçalves RL, Ramos IB, Koerich LB, et al. Silencing of maternal heme-binding protein causes embryonic mitochondrial dysfunction and impairs embryogenesis in the blood sucking insect Rhodnius prolixus. Journal of Biological Chemistry. 2013;288: 29323–29332.

33. Merril CR, Dunau ML, Goldman D. A Rapid Sensitive Silver Stain for Polypeptides in Polyacrylamide Gels. Anal Biochem. 1981;110: 201–207.

34. Lowry OH, Rosebrough NJ, Farr AL, Randall RJ. PROTEIN MEASUREMENT WITH THE FOLIN PHENOL REAGENT. J Biol Chem. 1951;193: 265–275.

35. Oliveira PL, Kawooya JK, Ribeiro JM, Meyer T, Poorman R, Alves EW, et al. A heme-binding protein from hemolymph and oocytes of the blood-sucking insect, Rhodnius prolixus. Isolation and characterization. J Biol Chem. 1995;270: 10897–10901.

36. Liu QH, Zhang SC, Li ZJ, Gao CR. Characterization of a pattern recognition molecule vitellogenin from carp (Cyprinus carpio). Immunobiology. 2009;214: 257–267. doi:10.1016/j.imbio.2008.10.003

37. Nunes DLM, Carvalho-Araujo MF, Silva-Cabral S, Rios T, Chagas-Lima AC, de Sousa G, et al. Lipid metabolism dynamic in Triatomine Rhodnius prolixus during acute Trypanosoma rangeli infection. Acta Trop. 2023;248. doi:10.1016/j.actatropica.2023.107032

38. Fan Y, Schal C, Vargo EL, Bagnères A-G. Characterization of termite lipophorin and its involvement in hydrocarbon transport. J Insect Physiol. 2004;50: 609–620.

39. Majerowicz D, Cezimbra MP, Alves-Bezerra M, Entringer PF, Atella GC, Sola-Penna M, et al. Rhodnius Prolixus Lipophorin: Lipid Composition And Effect Of High Temperature On Physiological Role. Arch Insect Biochem Physiol. 2013;82: 129–140. doi:10.1002/arch.21080

40. Ramos IB, Miranda K, de Souza W, Machado EA. Calcium-regulated fusion of yolk granules during early embryogenesis of Periplaneta americana. Mol Reprod Dev. 2006;73: 1247–1254.

41. Ramos IB, Miranda K, de Souza W, Oliveira DMP, Lima a PC a, Sorgine MHF, et al. Calcium-regulated fusion of yolk granules is important for yolk degradation during early embryogenesis of Rhodnius prolixus Stahl. J Exp Biol. 2007;210: 138–48. doi:10.1242/jeb.02652

42. Macedo-Silva A, Rios T, Ramos I, Majerowicz D. Lipophorin receptor knockdown reduces hatchability of kissing bug Rhodnius prolixus eggs. Insect Biochem Mol Biol. 2025;176: 104221. 10.1016/j.ibmb.2024.104221

43. Leyria J. Endocrine factors modulating vitellogenesis and oogenesis in insects: An update. Mol Cell Endocrinol. 2024; 112211.

44. Leyria J, Orchard I, Lange AB. Impact of JH Signaling on Reproductive Physiology of the Classical Insect Model, Rhodnius prolixus. International Journal of Molecular Sciences. 2022. doi:10.3390/ijms232213832

45. Benrabaa SAM, Orchard I, Lange AB. The role of ecdysteroid in the regulation of ovarian growth and oocyte maturation in Rhodnius prolixus, a vector of Chagas disease. Journal of Experimental Biology. 2022;225: jeb244830.

46. Benrabaa S, Orchard I, Lange AB. A critical role for ecdysone response genes in regulating egg production in adult female Rhodnius prolixus. PLoS One. 2023;18: e0283286.

47. Leyria J, Benrabaa S, Nouzova M, Noriega FG, Tose LV, Fernandez-Lima F, et al. Crosstalk between nutrition, insulin, juvenile hormone, and ecdysteroid signaling in the classical insect model, Rhodnius prolixus. Int J Mol Sci. 2022;24: 7.

48. Gondim K, Oliveira P, Coelho H, Masuda H. Lipophorin from Rhodnius prolixus: purification and partial characterization. Insect Biochem. 1989;19: 153–161.

49. Gondim KC, Oliveira PL, Masuda H. Lipophorin and oögenesis in Rhodnius prolixus: Transfer of phospholipids. J Insect Physiol. 1989;35: 19–27. 10.1016/0022-1910(89)90032-2

50. Faria FS, Garcia ES, Goldenberg S. Synthesis of a haemolymph hexamerin by the fat body and testis of Rhodnius prolixus. Insect Biochem Mol Biol. 1994;24: 59–67.

51. Paiva-Silva GO, Sorgine MHF, Benedetti CE, Meneghini R, Almeida IC, Machado EA, et al. On the biosynthesis of Rhodnius prolixus heme-binding protein. Insect Biochem Mol Biol. 2002;32: 1533–1541. 10.1016/S0965-1748(02)00074-7

52. Machado EA, Oliveira PL, Moreira MF, De Souza W, Masuda H. Uptake of Rhodnius heme-binding protein (RHBP) by the ovary of Rhodnius prolixus. Arch Insect Biochem Physiol. 1998. doi:10.1002/(sici)1520-6327(1998)39:4<133::aid-arch1>3.0.co;2-d

53. Vieira PH, Bomfim L, Atella GC, Masuda H, Ramos I. Silencing of RpATG6 impaired the yolk accumulation and the biogenesis of the yolk organelles in the insect vector R. prolixus. PLoS Negl Trop Dis. 2018;12: e0006507. doi:10.1371/journal.pntd.0006507

54. de Almeida E, Dittz U, Pereira J, Walter-Nuno AB, Paiva-Silva GO, Lacerda-Abreu MA, et al. Functional characterization of maternally accumulated hydrolases in the mature oocytes of the vector Rhodnius prolixus reveals a new protein phosphatase essential for the activation of the yolk mobilization and embryo development. Front Physiol. 2023;14. doi:10.3389/fphys.2023.1142433

55. Vieira PH, Benjamim CF, Atella G, Ramos I. VPS38/UVRAG and ATG14, the variant regulatory subunits of the ATG6/Beclin1-PI3K complexes, are crucial for the biogenesis of the yolk organelles and are transcriptionally regulated in the oocytes of the vector Rhodnius prolixus. PLoS Negl Trop Dis. 2021;15. doi:10.1371/journal.pntd.0009760

56. Mitchell RD, Sonenshine DE, Pérez De León AA. Vitellogenin receptor as a target for tick control: A mini-review. Frontiers in Physiology. Frontiers Media S.A.; 2019. doi:10.3389/fphys.2019.00618

57. Huo Y, Liu W, Zhang F, Chen X, Li L, Liu Q, et al. Transovarial Transmission of a Plant Virus Is Mediated by Vitellogenin of Its Insect Vector. PLoS Pathog. 2014;10. doi:10.1371/journal.ppat.1003949

58. Mao Q, Wu W, Huang L, Yi G, Jia D, Chen Q, et al. Insect Bacterial Symbiont-Mediated Vitellogenin Uptake into Oocytes To Support Egg Development. 2020. doi:10.1128/mBio

59. Freitak D, Schmidtberg H, Dickel F, Lochnit G, Vogel H, Vilcinskas A. The maternal transfer of bacteria can mediate trans-generational immune priming in insects. Virulence. 2014;5: 547–554. doi:10.4161/viru.28367

60. Zhang S, Wang S, Li H, Li L. Vitellogenin, a multivalent sensor and an antimicrobial effector. International Journal of Biochemistry and Cell Biology. Elsevier Ltd; 2011. pp. 303–305. doi:10.1016/j.biocel.2010.11.003

61. Morini M, Lafont AG, Maugars G, Baloche S, Dufour S, Asturiano JF, et al. Identification and stable expression of vitellogenin receptor through vitellogenesis in the European eel. Animal. 2020;14: 1213–1222. doi:10.1017/S1751731119003355

62. Okabayashi K, Shoji H, Nakamura T, Hashimoto O, Asashima M, Sugino H. cDNA Cloning and Expression of theXenopus laevisVitellogenin Receptor. Biochem Biophys Res Commun. 1996;224: 406–413.

63. Bidwell CA, Carlson DM. Characterization of vitellogenin from white sturgeon, Acipenser transmontanus. J Mol Evol. 1995;41: 104–112.

64. Liu X, Liu C, Feng Y, Guo X, Zhang L, Wang M, et al. Male vitellogenin regulates gametogenesis through a testis-enriched big protein in Chrysopa pallens. Insect Mol Biol. 2024;33: 17–28.

65. Lidani KCF, Andrade FA, Bavia L, Damasceno FS, Beltrame MH, Messias-Reason IJ, et al. Chagas disease: From discovery to a worldwide health problem. Journal of Physical Oceanography. American Meteorological Society; 2019. doi:10.3389/fpubh.2019.00166

66. Gómez-Ochoa SA, Rojas LZ, Echeverría LE, Muka T, Franco OH. Global, Regional, and National Trends of Chagas Disease from 1990 to 2019: Comprehensive Analysis of the Global Burden of Disease Study. Glob Heart. 2022;17. doi:10.5334/GH.1150

67. Ribeiro JMC, Genta FA, Sorgine MHF, Logullo R, Mesquita RD, Paiva-Silva GO, et al. An Insight into the Transcriptome of the Digestive Tract of the Bloodsucking Bug, Rhodnius prolixus. PLoS Negl Trop Dis. 2014;8: 27. doi:10.1371/journal.pntd.0002594

68. Medeiros MN, Logullo R, Ramos IB, Sorgine MHF, Paiva-Silva GO, Mesquita RD, et al. Transcriptome and gene expression profile of ovarian follicle tissue of the triatomine bug rhodnius prolixus. Insect Biochem Mol Biol. 2011;41: 823–831. doi:10.1016/j.ibmb.2011.06.004

69. Coelho VL, de Brito TF, de Abreu Brito IA, Cardoso MA, Berni MA, Araujo HMM, et al. Analysis of ovarian transcriptomes reveals thousands of novel genes in the insect vector Rhodnius prolixus. Sci Rep. 2021;11. doi:10.1038/s41598-021-81387-1

70. Leyria J, Orchard I, Lange AB. Transcriptomic analysis of regulatory pathways involved in female reproductive physiology of Rhodnius prolixus under different nutritional states. Sci Rep. 2020;10: 1–16.

71. Pereira J, Rios T, Amorim J, Faria-Reis A, de Almeida E, Neves M, et al. Functional characterization of vitellogenin unveils novel roles in RHBP uptake and lifespan regulation in the insect vector Rhodnius prolixus. Insect Biochem Mol Biol. 2025;180: 104301.

72. Yoth M, Maupetit-Méhouas S, Akkouche A, Gueguen N, Bertin B, Jensen S, et al. Reactivation of a somatic errantivirus and germline invasion in Drosophila ovaries. Nat Commun. 2023;14. doi:10.1038/s41467-023-41733-5

73. Brasset E, Taddei AR, Arnaud F, Faye B, Fausto AM, Mazzini M, et al. Viral particles of the endogenous retrovirus ZAM from Drosophila melanogaster use a pre-existing endosome/exosome pathway for transfer to the oocyte. Retrovirology. 2006;3: 25.

74. Jia D, Mao Q, Chen Y, Liu Y, Chen Q, Wu W, et al. Insect symbiotic bacteria harbour viral pathogens for transovarial transmission. Nat Microbiol. 2017;2: 1–7.

75. Wei J, He Y-Z, Guo Q, Guo T, Liu Y-Q, Zhou X-P, et al. Vector development and vitellogenin determine the transovarial transmission of begomoviruses. Proceedings of the National Academy of Sciences. 2017;114: 6746–6751.

76. Guo Q, Shu Y-N, Liu C, Chi Y, Liu Y-Q, Wang X-W. Transovarial transmission of tomato yellow leaf curl virus by seven species of the Bemisia tabaci complex indigenous to China: Not all whiteflies are the same. Virology. 2019;531: 240–247.

77. Salmela H, Amdam G V, Freitak D. Transfer of immunity from mother to offspring is mediated via egg-yolk protein vitellogenin. PLoS Pathog. 2015;11: e1005015.

